# Ultrafast end-to-end protein structure prediction enables high-throughput exploration of uncharacterised proteins

**DOI:** 10.1101/2020.11.27.401232

**Authors:** Shaun M Kandathil, Joe G Greener, Andy M Lau, David T Jones

## Abstract

Deep learning-based prediction of protein structure usually begins by constructing a multiple sequence alignment (MSA) containing homologues of the target protein. The most successful approaches combine large feature sets derived from MSAs, and considerable computational effort is spent deriving these input features. We present a method that greatly reduces the amount of preprocessing required for a target MSA, while producing main chain coordinates as a direct output of a deep neural network. The network makes use of just three recurrent networks and a stack of residual convolutional layers, making the predictor very fast to run, and easy to install and use. Our approach constructs a directly learned representation of the sequences in an MSA, starting from a one-hot encoding of the sequences. When supplemented with an approximate precision matrix, the learned representation can be used to produce structural models of comparable or greater accuracy as compared to our original DMPfold method, while requiring less than a second to produce a typical model. This level of accuracy and speed allows very large-scale 3-D modelling of proteins on minimal hardware, and we demonstrate that by producing models for over 1.3 million uncharacterized regions of proteins extracted from the BFD sequence clusters. After constructing an initial set of approximate models, we select a confident subset of over 30,000 models for further refinement and analysis, revealing putative novel protein folds. We also provide updated models for over 5,000 Pfam families studied in the original DMPfold paper.

**Significance Statement:** We present a deep learning-based predictor of protein tertiary structure that uses only a multiple sequence alignment (MSA) as input. To date, most emphasis has been on the accuracy of such deep learning methods, but here we show that accurate structure prediction is also possible in very short timeframes (a few hundred milliseconds). In our method, the backbone coordinates of the target protein are output directly from the neural network, which makes the predictor extremely fast. As a demonstration, we generated over 1.3 million models of uncharacterised proteins in the BFD, a large sequence database including many metagenomic sequences. Our results showcase the utility of ultrafast and accurate tertiary structure prediction in rapidly exploring the “dark space” of proteins.

## 1. Introduction

Analysis of amino acid residue covariation in deep multiple sequence alignments (MSAs) has revealed that covarying residues in proteins are frequently found to be close together in the tertiary structure. This principle has been successfully exploited for predicting inter-residue contacts by a variety of methods. Notable among these are the family of methods based on direct-coupling analysis (DCA) (1–3), and more recently, a number of methods based on deep learning. The latter have produced increasingly precise predictions of inter-residue contacts in recent years (4–6), and have also been adapted to output probabilistic predictions of the *distance* between the residues in contact, either as a probability distribution (7–10), or as real values (11–13). These predictions are usually made by models that operate on precomputed features derived from an MSA, such as covariance and/or precision matrices, contact predictions from DCA-based methods, together with other features such as predicted secondary structure labels and sequence profiles. This approach, though effective, requires that these features be precomputed, which can in some cases be a time-consuming process and can sometimes take longer than the rest of the prediction pipeline combined. Additionally, it does not allow the neural network (NN) model to utilise all the information that might be available in the sequence alignment, as a model trained on derived features is limited to using the information available in those handpicked features.

A key difficulty in using MSAs directly as input to a neural network is the fact that an MSA for a target sequence can have arbitrarily many sequences in it. To date, only a few published methods attempt to use an encoding of the MSA itself as the input to a neural network-based predictor of protein structural features. In rawMSA (14), the difficulty of embedding MSAs of arbitrary depth was addressed by training separate convolutional networks using predetermined maximum MSA depths. Recently, we demonstrated that a system of recurrent neural network (RNN) layers can be used to process MSAs of arbitrary size (15). A handful of subsequent studies have also demonstrated that directly processing MSAs in a neural network can be effective for predicting structural features such as inter-residue contacts or distances. In the MSA Transformer (16), the neural network model exploits the equivalence of aligned positions in the MSA in order to derive embeddings via a tied row attention framework. These embeddings are shown to be useful for producing accurate predictions of inter-residue contacts in a number of proteins.The process of end-to-end differentiable coordinate generation for protein structures, where atomic coordinates are produced as a direct output of a neural network, was first demonstrated in the Recurrent Geometric Network (18), which takes as input an amino acid sequence and a position-specific substitution matrix (PSSM) for the sequence family. Another end-to-end method, NEMO (19), combines a neural-network representation of a forcefield with a coarse-grained Langevin dynamics simulator to derive structures, starting from a sequence and PSSM.

A number of studies have applied recent advances in protein structure prediction to model large sets of protein sequences with unknown structures, such as Pfam families (9, 20–23) or proteins from a minimal genome (24). These efforts have been limited, though, by the computational cost of running more than a few thousand predictions in a reasonable time frame. This has prevented structural annotations from keeping pace with the vast increase in sequence data now available as a result of metagenomic sequencing efforts. These large sequence collections present unique opportunities for discovery of proteins with potentially novel folds and/or novel functionality, and therefore, the development of rapid, accurate methods for predicting protein structures from sequence information alone is highly desirable.

In this paper, we develop a deep neural network-based method for predicting protein structures directly from MSAs in an end-to-end-differentiable fashion, building on methods we trialled in CASP14 (15). Our method produces coordinates of backbone atoms as a direct output of the neural network model, further saving time over model generation approaches used previously, and also provides a predicted confidence in the output coordinates. Using this method, we model protein sequences from the BFD (25, 26), a clustered collection of over 2.2 billion sequences assembled from a number of metagenomic sequencing efforts as well as sequences from UniProt. To our knowledge, no study has attempted to leverage this collection of sequences to derive structural annotations for the protein families represented therein.

We show that despite the relatively simple composition of the input feature set, we are able to obtain predictions of comparable or better accuracy than DMPfold (9), while taking under a second to a few minutes to produce one structure on readily-available hardware. DMPfold2 is thus fast enough to run at unprecedented scale, and we apply it to over 1.3 million subsequences without detectable templates extracted from the BFD, generating over 30,000 confident models, and using these to explore the structural content of the dark protein space in the BFD. We show how such an approach can be used to find remote homologues and aid in function prediction, along with finding a number of novel folds. We also provide updated models for Pfam families whose structures were predicted in the original DMPfold study (9).

## 2. Materials and Methods

### 2.1. Datasets for training and evaluation

Training was conducted on a mixture of both chains and domains, comprising 31,159 domains from the V4.2 CATH (27) s35 representative set of domains and 6,742 full length chains from the original DMPfold1 training set. A set of 300 chains were held out from training to use as a validation set to monitor convergence. MSAs for each training example in the CATH s35 set were created using 3 iterations of HHblits v3.0b3 (28) using the UniClust30 (October 2017) database, with an E-value threshold of 0.001 and minimum coverage of 50%.

We tested the effectiveness of our method on a set of 39 domains from the CASP13 experiment, categorized as either FM or FM/TBM by the CASP13 assessors. MSAs were built using HHblits (28) by first searching the UniRef30 database (2020_06 release), then using the MSA thus obtained as a query for a further HHblits search against the BFD.

### 2.2. Neural Network Model architecture, input and output features

A schematic representation of the DMPfold2 architecture is shown in Figure 1a, and the MSA embedding procedure is shown in Figure 1b. We used a system of gated recurrent unit (GRU) layers to process and embed the input MSA, and output a feature map with 256 x L values, where L is the length of the target sequence. Starting from a 22-dimensional one-hot encoding of each residue in each sequence in the MSA, a stack of 2 GRU layers scans individual columns in the MSA in the vertical direction. The hidden state of these layers is a vector of fixed size (512 in our case), and the hidden state at the end of the vertical scan is used as a fixed-length representation of the information in each column of the MSA. The per-column representations are then passed to another stack of 2 bidirectional GRU layers with 256 hidden units, producing final embeddings for each column of 2 x 256 values. These embeddings are striped vertically and horizontally (similar to the manner in which per-residue features are prepared for use by 2D convolutional layers (4, 5)), and combined with a fast approximation of a precision matrix, calculated using the “fast_dca” algorithm from trRosetta (10). These features are then used as input to a convolutional maxout layer to reduce the number of channels to 128, followed by a series of 16 residual blocks, each composed of one maxout layer (29) and one squeeze-excitation layer (30) each. As we first showed with DMPfold1 (9), iteration of the prediction process, where the final model is fed back in as an input to the network, can make significant improvements in model quality. DMPfold2 also allows iterations, where the output coordinate set can be used as a seed for second and subsequent passes through the network. To allow this, the ResNet incorporates an extra input feature channel carrying pairwise Cα distances from an existing 3-D structure, which can be set to a constant of -1 to indicate no distances are to be considered i.e. during the first pass.

**Figure 1.**
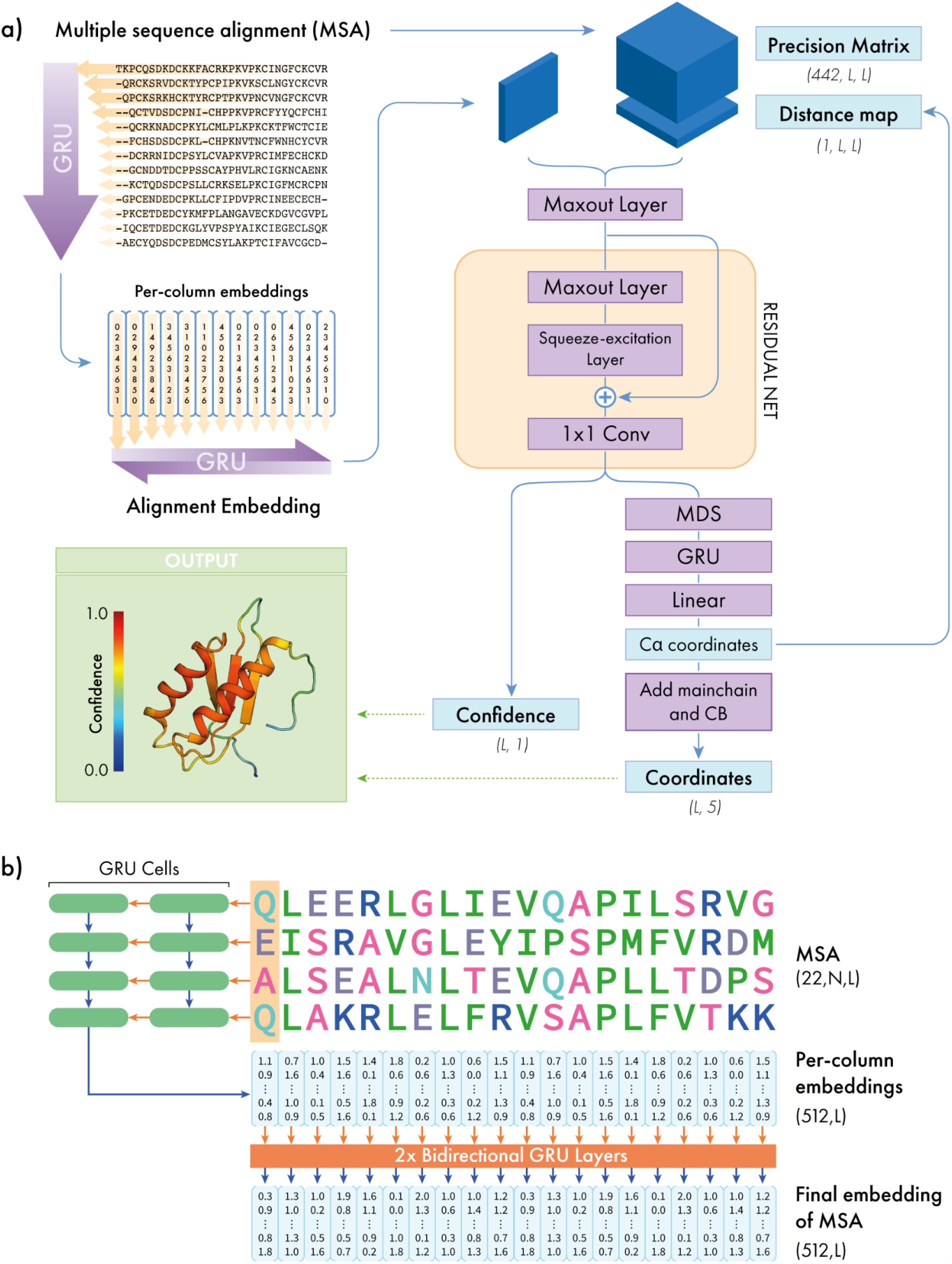
Schematic representation of the DMPfold2 neural net. **a)** The input multiple sequence alignment (MSA) is processed into a learned embedding, as shown in detail in b). This embedding is combined with a precision matrix calculated from the MSA, and fed to a convolutional Maxout layer to reduce its dimensionality, before being fed to a series of 16 residual neural net (ResNet) blocks. Each block is composed of a convolutional Maxout layer and a Squeeze-excitation layer. The outputs from the network are collected as the outputs of a 2D convolutional layer with 1×1 filter, and are a combination of different structural features, represented in the right-hand column. All outputs from the network are predicted jointly. **b)** Details of the MSA embedding section. The MSA is represented by a one-hot encoding of 22 residue types (including gaps and unknown residues). First, the residues in a single column of the MSA are treated as timesteps and fed as input to a stack of two Gated Recurrent Unit (GRU) network layers. The final hidden state of the second GRU, obtained after processing the whole column of the MSA, is used as an embedding of the information in that column. The process is then repeated for the remaining columns in the MSA, producing a separate embedding for each MSA column. Finally, these per-column embeddings are used as inputs to a stack of 2 bidirectional GRU layers that produces an embedding of all sequences and columns in the MSA. The dimensions of the input tensor and embeddings are shown in parentheses.

The final stages of the network implement end-to-end prediction of coordinates from the output embeddings of the ResNet as shown in Figure 1a. A two-channel 1×1 convolutional layer is used to project the ResNet embeddings into two channels, one for the output Cα-Cα distance map, the other to represent confidence estimates for each pair distance. The distance outputs from this layer are converted to a real-value distance matrix *D*by first averaging the upper and lower triangles of the matrix and taking absolute values to ensure that the distance matrix is positive-definite and symmetric. This matrix is projected to 3-D Cα coordinates using multi-dimensional scaling (31). The following (Gram) matrix is defined:

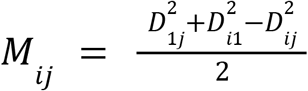

The eigendecomposition *M* = *USU*^*T*^ is then calculated, where 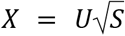 gives the coordinates of the points. Note that if a distance map can be fully embedded in 3-D space there will only be 3 non-zero eigenvalues of M, but this is not necessarily the case for the real-valued distance matrix predicted by the network. Coordinates corresponding to the largest 8 eigenvalues were therefore calculated and the rest discarded. One problem with MDS is that there are no learnable elements, and so there is no way for the network to learn to modify the coordinates to, for example, correct chirality issues. Therefore, a “learnable MDS” module was implemented simply by feeding the 8-dimensional output from the MDS step, concatenated with the original biGRU embedding of the MSA, through a final 2-layer bidirectional GRU recurrent network (256 weights per hidden layer), with 520 input channels and a final fully connected layer with 512 inputs and 3 outputs (representing the final Cα coordinates). During training with a rigid body coordinate loss, as opposed to a distance-based loss, this final network is able to learn to automatically transform the output coordinates for any structures mirrored by the multidimensional scaling process, without manual intervention.

After the network predicts Cα coordinates, a simple coordinate refinement module attempts to correct “obvious” issues with the model. Cα atoms adjacent in sequence are moved to a target distance of 3.78 Å and all Cα-Cα distances are moved to a minimum of 3.0 Å according to a harmonic potential for 100 steps. Backbone N, C, and O atoms and Cβ atoms are then added using a geometric procedure based on the “catomain” routine from the DRAGON method (32). Suppose that ***v***_**-1**_ is the vector from a Cα to the previous Cα in the sequence, ***v***_**+1**_ is the vector from a Cα to the next Cα in the sequence, ***m***_**-1**_ is the midpoint between a Cα and the previous Cα in the sequence, and ***x***_**α**_ is the normalised vector cross product of ***v***_**-1**_ and ***v***_**+1**_. The coordinates ***C***_**N**_ of N, ***C***_**C**_ of the previous residue C and ***C***_**O**_ of the previous residue O are then given by:

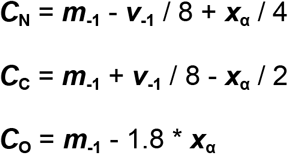

If ***v***_**N**_ is the vector from N to Cα in a residue, ***v***_**C**_ is the vector from C to Cα in a residue, ***v***_**β**_ is the normalised sum of ***v***_**N**_ and ***v***_**C**_, ***x***_**β**_ is the normalised vector cross product of ***v***_**N**_ and ***v***_**C**_, and ***C***_**α**_ is the coordinates of Cα, then the coordinates ***C***_**β**_ of Cβ are given by:

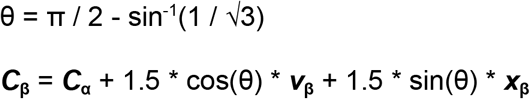

Special considerations are required for the terminal residues, with details available in the source code. No further atom-level optimisation or refinement of the structure was attempted in any of the results shown, though it would be easy to add side chains to these models and refine the models further, at the expense of losing the end-to-end characteristics of the method.

In its iterative mode, the DMPfold2 network allows a recurrent processing of the final model. In this mode, predicted Cα coordinates are used to calculate a distance matrix and this matrix is fed back to the ResNet layer as an additional input channel. During training, the number of allowed iterations was varied between 0 and 5 due to the accumulated gradients and GPU memory limitations, but during inference any number of iterations can be requested as no gradient information is needed. The final model returned after iteration is the model which has the highest overall confidence as judged by one output channel of the ResNet.

### 2.3. Training procedures

The DMPfold2 neural network was implemented using PyTorch (33) and trained using the Adam optimizer (34). An initial learning rate of 0.0003 was used, a dropout probability of 0.1 was used for the recurrent network layers, and dropout 0.2 used in the convolutional layers. For data augmentation, at each epoch each alignment in the training set is subsampled as follows:

a. Random rows from the alignment are selected, up to a maximum of 1,000, though always including the target sequence in the first row.
b. Alignments with > 350 columns (residues) are randomly cropped to that maximum length to fit within available GPU memory, and the target coordinates are similarly cropped to match.

As a loss function, we opted for TM-score (35) rather than RMSD. This choice of loss function reduced the effects of bad model outliers and kept both the model loss and confidence scores to be normalized to the range [0,1]. Confidence values are output per residue, which allows confidently modelled segments of the model to be distinguished from less confidently modelled segments.

### 2.4. Modelling protein domains from the BFD

#### 2.4.1. Extraction of sub-alignments from the BFD

The consensus sequences from each MSA in the BFD were extracted and scanned against the Pfam (36) and CATH-Gene3D (27, 37) HMM libraries using hmmscan (38) in order to filter out regions corresponding to well-characterised protein families. Unmatched subsequences at least 30 residues long were then used to extract the corresponding (sub-)MSAs from the BFD alignments, discarding rows in the resulting sub-alignment comprising only gap characters. The resulting set of 1,333,242 MSAs were used as the inputs for the modelling runs.

#### 2.4.2. Generation and evaluation of end-to-end-predicted models

We ran DMP2 using one iteration in order to obtain a first set of models. Because the BFD sub-alignments vary greatly in terms of quality and coverage, we opted to use a quick modelling sweep of the MSAs in order to identify promising candidates that might benefit from several iterations of the modelling procedure. In total, we generated 1,333,242 models from the BFD alignments by running one iteration of DMPfold2. From each model, we recorded model descriptors including: the number of target residues (nres), alignment depth, predicted confidence, radius of gyration (Rg) and 8-class secondary structure assignments via DSSP (39).

Of the 1,333,242 models, 48,625 models (3.65%) corresponded to single- or double-helix structures. We identified these models using the criteria that such models must consist of only one or two helices (composed of at least 50% of residues) and have a Rg/nres ratio of greater than 0.28 or 0.18, respectively. These criteria were manually calibrated from a sample of generated models. The Rg/nres ratio describes the average contribution of each residue towards the model Rg, and can be helpful for identifying single/double helices with partially malformed secondary structure.

Next, from the remaining 1,284,617 models, 33,548 (2.61%) were found to have a network-predicted confidence score of 0.4 or greater. These targets were scanned for disorder using IUPred (40, 41). To assess the similarity of these models to known folds, we compared each model to a representative set of PDB chains formed by clustering all protein chains in the PDB with 30+ residues at 30% sequence identity. The closest match by TM-score calculated using TM-align (42) normalised over the length of the modelled protein was recorded, meaning that models could register a match to part of a larger domain. To assign CATH superfamilies, any CATH domains present in the closest matched PDB chains were extracted using BioStructures.jl (43) and the TM-score was calculated with TM-align between the modelled protein and the CATH domain. The model was assigned to the superfamily if the largest TM-score was greater than 0.5. A visual summary of the entire model filtering workflow can be found in Supplementary Figure S1.

#### 2.4.3. Iterative refinement of medium-to-high-confidence models

225 models with a variety of predicted confidence scores were taken from the initial set of 1.3M models. We rebuilt MSAs for this set using HHblits searching against the BFD and modelled them using either 1 or 10 iterations of DMPfold2 modelling. For a baseline, we also used the same MSAs used in the first modelling sweep with either 1 or 10 iterations of DMPfold2 to evaluate the effect of rebuilding MSAs and including more iterations during modelling. We assessed the refined models in terms of their predicted confidence scores, secondary structure content, and similarity to PDB entries. Secondary structure content was assessed as the fraction of residues having a DSSP-assigned code corresponding to secondary structures. Models were compared to a representative set of PDB chains as described previously and potential novel folds were identified as those with a largest TM-score of less than 0.5.

### 2.5. Modelling Pfam families

In the original DMPfold1 publication (9), we generated models for a set of 5,214 Pfam families for which no structural annotation was available. Models were generated for 5,193 of these families, using DMPfold2 and the same alignments as those used for DMPfold1. Some target sequences contained non-standard amino acids and were excluded.

Experimental structures have since been made available for 255 of these families. To assess the accuracy of our models, we calculated the TM-score between the 255 newly annotated Pfam families and their respective PDB representatives. We additionally compared the accuracy of our models with those produced by DMPfold1 and C-I-TASSER (23) (Figure 2c-d). In this comparison, all generated models were compared to the same reference PDB chain via TM-align. Targets were additionally subdivided according to their predicted difficulties reported in C-I-TASSER (23). Overall, 231 and 255 models were compared between DMPfold2 vs DMPfold1, and DMPfold2 vs C-I-TASSER, respectively. In all cases, TM-scores were calculated via TM-align and were normalised by the length of the target protein. It is also important to note that the representative PDBs of the 255 families did not overlap with the DMPfold2 training set.

**Figure 2.**
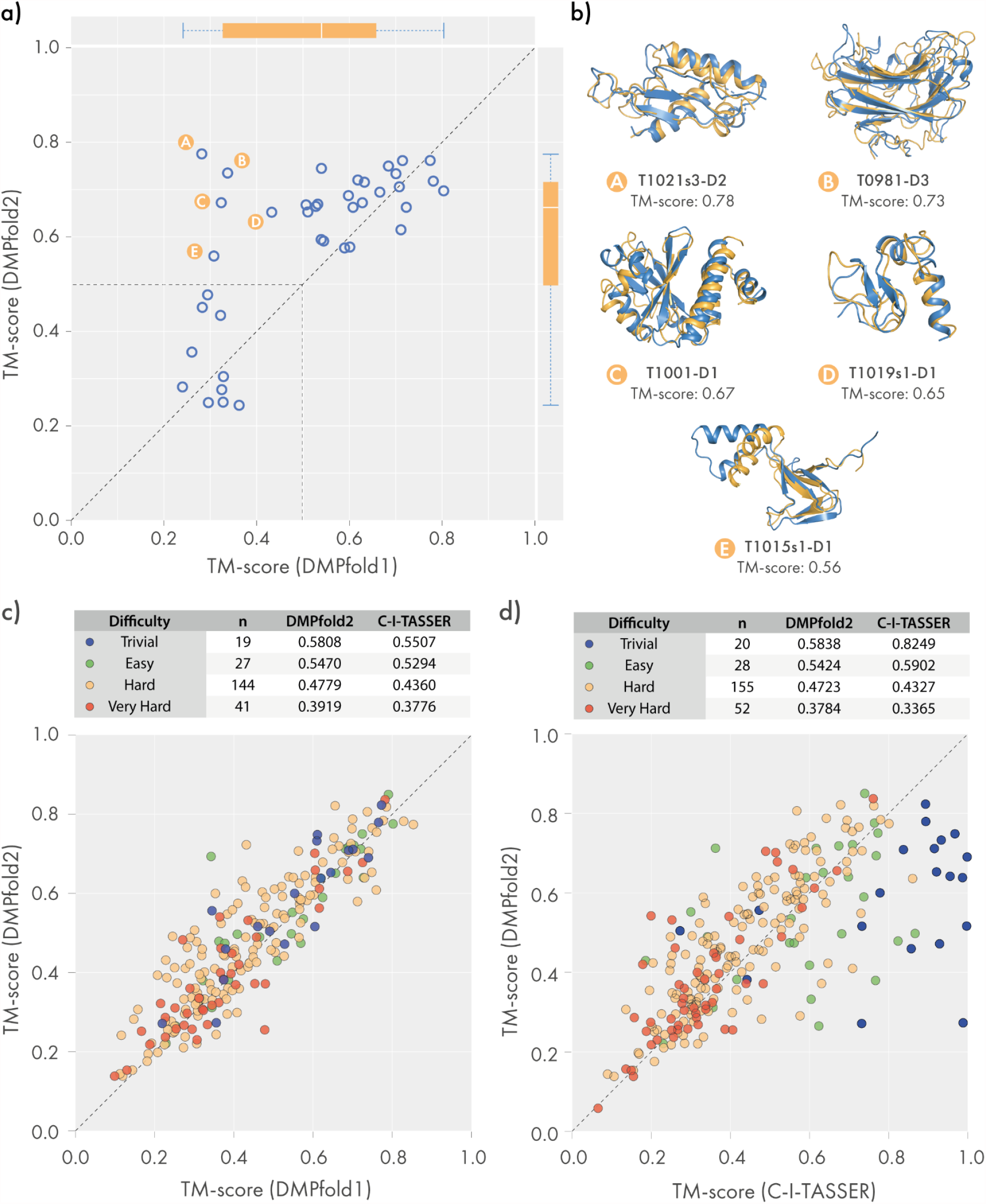
TM-scores obtained on CASP13 domains and Pfam families. **a**) Comparison of TM-scores obtained by DMPfold2 and DMPfold1 on 39 CASP FM and FM/TBM domains. Detailed model accuracy data for each target can be found in Table S1. A dashed line of unit slope is drawn, as well as segments demarcating the TM-score ≥ 0.5 regions of the plot. Overall, DMPfold2 end-to-end achieves comparable or greater model accuracy than DMPfold1. **b**) Five CASP13 domains could be folded correctly (TM-score > 0.5) by the end-to-end procedure but not by DMPfold1. Models and native structures are shown in orange and blue respectively. **c)** Comparison of Pfam families with recent structural annotations modelled by DMPfold2 vs DMPfold1 and **d**) DMPfold2 vs C-I-TASSER. The colour of each data point represents the target difficulty as reported in ref (23). The values in the tables represent the mean TM-score obtained by each method in each target category.

## 3. Results

### 3.1. Tertiary structure model accuracy on CASP13 domains

As an evaluation of the effectiveness of the DMPfold2 end-to-end model building procedure, we built 3-D models of 39 FM and FM/TBM domains from the CASP13 experiment. Comparison against models generated using DMPfold1 (Figure 2 and Table S1) shows that DMPfold2 produces more accurate structure models for these domains. DMPfold2 is able to fold 30 domains to a TM-score of 0.5 or greater, as compared to 26 for DMPfold1. The mean TM-score is 0.590 for DMPfold2 and 0.531 for DMPfold1.

### 3.2. Pfam models

We produced updated models for 5,193 Pfam families, using the same MSAs as used in the original DMPfold paper (9). Approximately 60% of the models generated were at a confidence score of 0.4 or greater, with the majority of these high-confidence models representing targets between 50-200 residues in length (Figure S4a-b).

255 of the modelled Pfam families have received mappings to PDB structures since the last DMPfold publication. As a benchmarking exercise, we compared the models of these 255 families generated by DMPfold2 to those generated by DMPfold1 and the recently published C-I-TASSER method (23), calculating the TM-score of each model to a PDB entry assigned to the equivalent family (Figure 2c-d). We additionally subdivided targets based on their LOMETS classification (23) into trivial, easy, hard and very hard categories. From our analysis, it can be seen that DMPfold2 outperforms DMPfold1 in all difficulty classes, in terms of the mean TM-score achieved for each category (Figure 2c). DMPfold2 further outperforms C-I-TASSER in the hard and very hard categories, but underperforms in trivial and easy cases. Examples of six targets belonging to the very hard category are shown in Figure S4c. The lower performance of our method on easy and trivial targets can be attributed to the fact that DMPfold2 does not make explicit use of template information, whereas C-I-TASSER does. The architecture of DMPfold2 does allow for distance information from templates to be added as an input to the ResNet section, replacing the distance map derived from the output coordinates (see Section 2.2), though this was not done during the training process. We are currently investigating methods for explicitly supplying template information during the prediction process.

### 3.3. BFD modelling experiments

We modelled 1,333,242 subsequences from the BFD not matching conserved families in the Pfam or CATH-Gene3D HMM libraries using one iteration of DMPfold2. Given that around 86% of these subsequences were fewer than 300 residues in length, most will correspond to single domains, and a single pass through the DMPfold2 neural networks will provide a rough idea of the distribution of folds present, following which we can select a subset of promising models for further refinement and analysis. A summary of key metrics on the set of 1.3 million models appears in Figure S2. Although the vast majority of the models are predicted to have a confidence score of less than 0.4, most of these low-confidence models correspond to target sequences that are either very short, and/or whose MSAs have very few sequences (Figure S2). Additionally, for some fraction of these targets, domain boundaries may not have been correctly parsed, as our subsequence filtering procedures were based only on sequence profile HMM similarity.

From the initial set of 1,333,242 models, we focused on a subset of 33,548 models predicted by DMPfold2 to have a confidence score of 0.4 or greater (see Methods). Our reasoning for not choosing a stricter cutoff was that it should be possible to refine some fraction of these models using the iterative prediction mode of DMPfold2. The majority of these targets are also predicted to be ordered according to IUPred, with 79% of sequences having 80% or more residues predicted as ordered. We assessed this set of models for similarity to existing CATH families. Overall, we were able to assign 23,495 models to 2,394 CATH superfamilies on the basis of structural similarity (Table S2), suggesting that the predictions are correct at the fold level. Figure 3 shows the distribution of these models and their folds. The large fraction of models with a similar structure in the PDB despite most sequences with templates being eliminated shows the value in *de novo* protein structure prediction for mapping sequences to the existing fold space and inferring homology. Indeed most targets in the FM category at CASP do match a known fold, even though the template is not found with sequence-based searching (44). As in our study of Pfam with DMPfold1, we do find a number of confident models that do not match structures in the PDB and likely represent novel folds.

**Figure 3.**
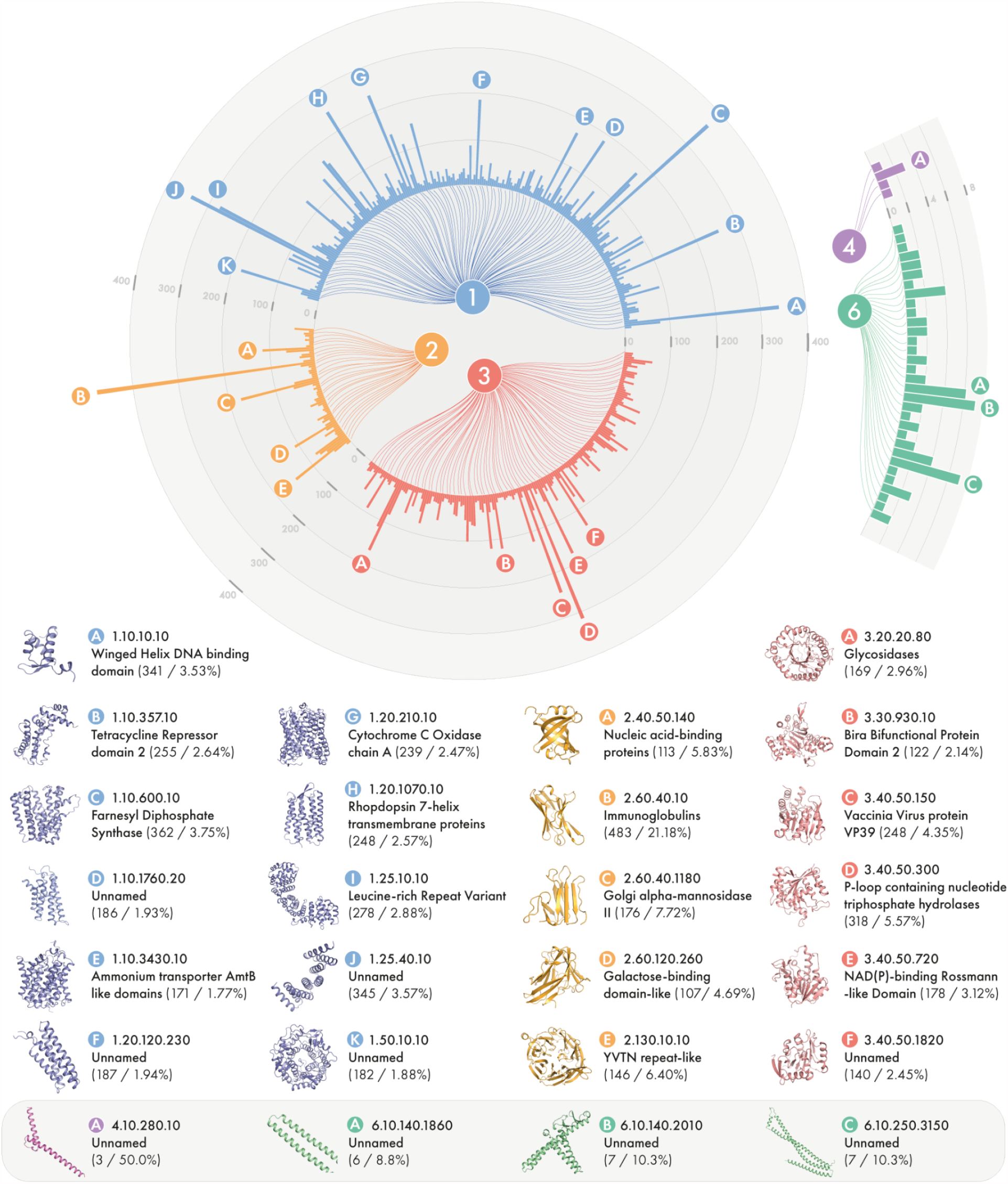
Distribution of BFD models assigned to CATH superfamilies. Data represents the number of models assigned to each class and superfamily from CATH. The values under each superfamily name represents the number of models assigned to that superfamily, and as a percentage of total models assigned to that class. Superfamilies with fewer than 10 models assigned for class 1-3 were omitted for clarity (211/945, 54/451 and 154/962 shown respectively).

#### 3.3.1. Iterative refinement of medium-to-high-confidence models

225 targets with medium-to-high confidence models were chosen and new MSAs and models were built for these targets by searching against the BFD. We compared the models built using the original MSAs extracted from the BFD against those built afresh using the trimmed consensus sequences as a query, with DMPfold2 running either 1 or 10 iterations. The results in Figures S3 demonstrate that regardless of the MSA used, the iterative updates to the coordinates output by DMPfold2 result in improved confidence scores, as well as greater secondary structure content. Concomitantly, a larger fraction of the models are found to have a hit to the PDB with a TM-score of 0.5 or greater. These results show the benefit of iterative structure refinement, but also demonstrate that model quality can usually be improved by generating a fresh MSA for the subsequence of interest, rather than merely extracting a region of the original BFD MSA for a longer sequence family. A number of targets were identified as having a potentially novel fold, based on comparison to the PDB with TM-align. Three of these models are shown in Figure 4a.

**Figure 4.**
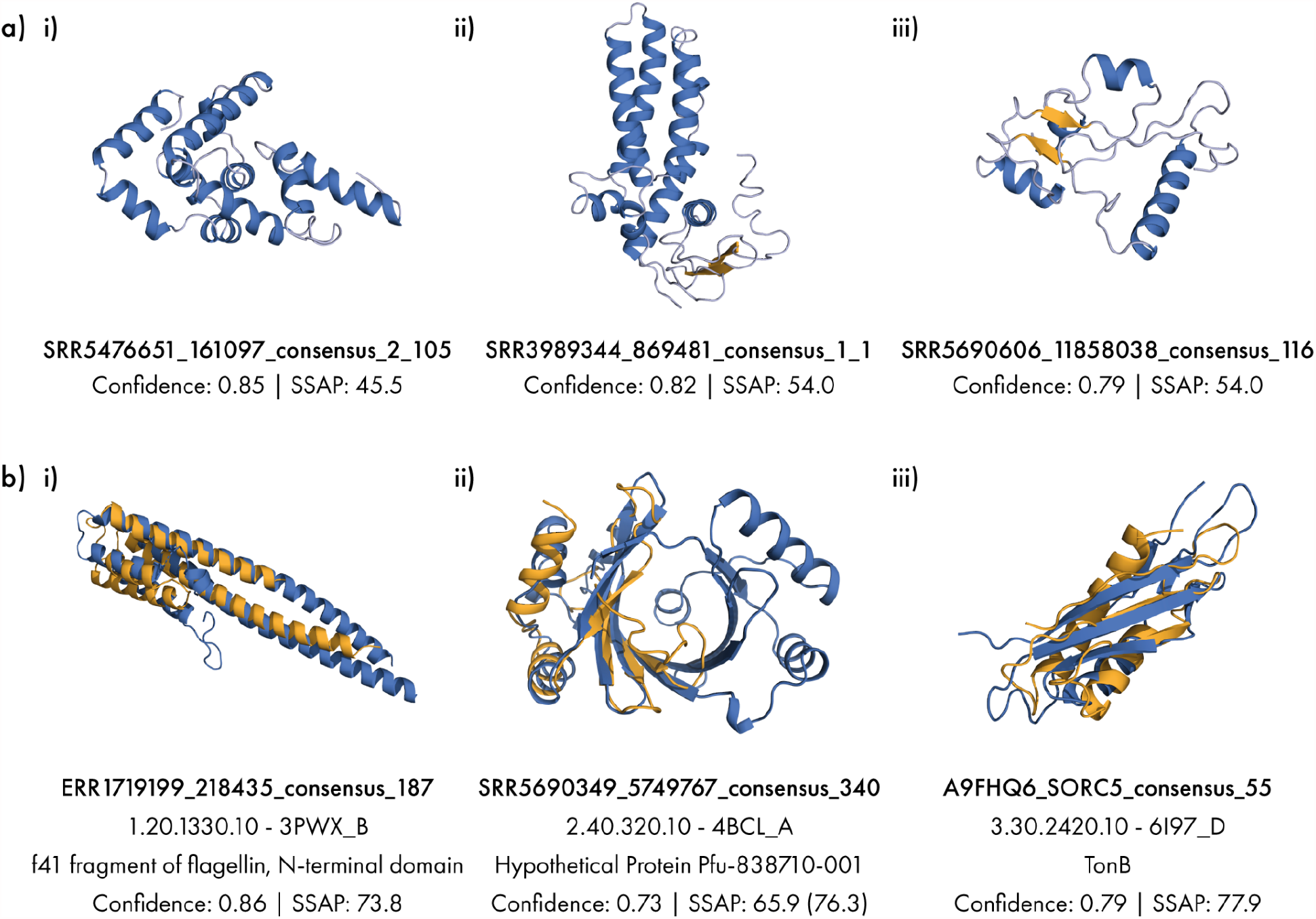
Examples of BFD models generated by DMPfold2. **a**) Potential novel folds identified from 225 remodelled BFD targets. Structures shown are high-confidence models with a SSAP score (55, 56) of less than 70 to the most similar structure in the PDB representative set. Models are generated from new BFD alignments using the 10-iteration end-to-end procedure. Helices and sheets are coloured in blue and orange respectively. **b**) Examples of BFD proteins matched to CATH topology classes with a single homologous superfamily. In these cases it is likely that the target protein shares function with other members of the superfamily. Models and native structures are shown in orange and blue respectively. For target b-ii), the value in parentheses represents the SSAP score if aligned to only half of the template.

#### 3.3.2. Template detection and function prediction using modelled structures

An additional experiment was run to assess whether the DMPfold2 models can be used for detection of distantly related proteins that may otherwise be missed by state of the art HMM-HMM comparison tools such as HHsearch. We used the BFD sub-MSAs for the 33,548 modelled sequences as queries to search the PDB70 database using HHsearch. 31% have at least one hit in the PDB70 database at an E-value threshold of 1.0. At an E-value threshold of 0.001, only 12% of the queries retrieve significant hits despite 83% having a model that matches a structure in the PDB. This illustrates the value of tertiary structure models in retrieving structural templates: assuming that the DMPfold2 models are correct at the fold level, the TM-align search against the PDB reveals that these models are structurally similar to existing PDB entries, whereas an HMM-HMM comparison would miss these templates. Of course, this principle has been well understood for some time, and is the basis of protein threading (45, 46), but the ease and speed with which a rough candidate model can be generated for a query sequence has the potential to accelerate and improve the sensitivity of structural template searches.

In some cases it is possible to speculate about the function of modelled sequences by matching the model to structures of known function (47). Three examples are shown in Figure 4b of models that match CATH superfamilies that are the only superfamily in the respective CATH topology type. In these cases it is likely that the sequences share a function with other members of the matched superfamily. This approach to assigning function will become more applicable as protein structure prediction methods improve and more experimental structure data becomes available.

## 4. Discussion

Recent successes in applying deep learning to protein structure prediction have mostly relied on the use of large, precomputed feature sets as inputs to the deep learning model. Here, we show that it is possible to directly process a MSA into a learned representation that is effective for predicting structural features in proteins. Structural models built using DMPfold2 are more accurate on average than those from DMPfold1. It is expected that allowing the network to access all the information in a raw MSA, rather than just pairwise frequency information, for example, enables it to extract richer information that can be used for more accurate prediction of structural features. Although the best performance is obtained when using the learned MSA representation alongside an on-the-fly computed precision matrix, the work in this study opens up the possibility of using just the MSA representation itself as the sole input to the ResNet for predicting protein structure, though that will probably require using methods with better ability to deal with very long range dependencies (see below), as some groups have recently done (16, 17). It also enables new lines of work that were prohibitive with large feature sets (such as those used in DMPfold1), due to the time and storage requirements of using those features.

The idea of end-to-end *de novo* prediction (18, 19) has certainly been tantalizing, in that a 3-D model can be produced in a fraction of a second compared to the hours or days needed previously. So far, however, most published end-to-end methods have not been able to produce models comparable to the state-of-the-art in *de novo* prediction, mainly because they have not effectively exploited covariation data as inputs. DMPfold2 is able to produce main chain coordinates of comparable or greater accuracy as compared to DMPfold1, while taking very little time to produce each model. This allows quick validation of *de novo* predictions before full modelling is carried out, or could be combined with a refinement method to almost completely replace the whole 3-D modelling pipeline. We showcased this approach by applying DMPfold2 to sequence families from the BFD, and even using a relatively coarse-grained approach, we were able to predict structures with plausible topologies for tens of thousands of uncharacterised protein domains, including some with potentially novel folds. We also showed that the modelling process can be used to identify fold-level similarity to existing PDB entries, where such similarity would otherwise have been missed by a purely sequence-based comparison.

During the preparation of our manuscript, two papers describing successful protein structure prediction using MSA embeddings were published. The RoseTTAFold method employs direct embeddings of MSAs to produce distance restraints, as well as to produce coordinates directly (17). This method employs a three-track attention-based neural network model inspired by DeepMind’s AlphaFold2 system, which produced excellent results in CASP14. AlphaFold2 (48) employs a number of novel neural network modules that are specialised for producing representations of MSAs and protein structures, beginning from a direct embedding of the input MSA and target sequence. In comparison to these methods, DMPfold2 produces models in much less time (Figure S5), making it suitable for use at a very large scale. Although DMPfold2’s models are not as accurate, they are still of sufficient accuracy to allow for easy identification of potentially interesting targets from very large datasets that can then be modelled using slower and more accurate modelling tools.

There are many interesting potential applications for nearly instant tertiary structure modelling methods based on direct embedding of MSAs. One example would be to visualize changes to the final 3-D model as the input sequences in the MSA are changed, essentially in real time. This opens up the possibility of *optimizing* the MSA in a way that maximises the neural network’s confidence in the model. Changes in predicted model confidence could also be used to identify profile drift in iterated HMM-HMM searches or clustering, which would in theory manifest as a drop in predicted model confidence upon incorporating additional sequence hits in the MSA. Predicted structures could also be used as enhancements to existing HMM libraries, similar to the manner in which secondary structure predictions have been used in the past (49). They could also be used to predict domain boundaries in large sequence libraries.

We are continuing to develop new neural network architectures for end-to-end structure prediction. In DMPfold2, the use of two gated recurrent networks to embed the input MSA is clearly effective, but was something of a design compromise in that RNNs are relatively fast to train, but have known limitations in terms of the limits of modelling long-range dependencies in sequences. In theory, gated RNNs are able to avoid the problems of vanishing gradients when modelling long sequences, but in practice, dependencies beyond a window of a few hundred time steps are poorly modelled. For this reason we used two GRU networks, one to embed in the vertical (sequence number) and one in the horizontal (residue number) direction so that the number of time steps that each GRU would need to model would be limited by either the lengths of typical protein domains or the depths of typical MSAs. For MSAs with longer sequences or with many more homologous sequences, even gated RNNs will start to become ineffective. We are currently investigating alternative means of embedding MSAs, such as the use of models based on new efficient transformer architectures (50), which are far more memory-efficient than the original Transformer (51) due to avoiding the calculation of large self-attention matrices over the length of the sequences. Standard transformer models have already been used to embed unaligned protein sequences (52–54), but the most efficient transformer models released in the last year are now capable of handling sequence lengths even in the millions, and so this suggests that a single deep transformer model with a compressed self-attention mechanism could, in principle, embed a whole MSA in one go by essentially treating it as a single sequence. Current experiments along these lines are promising.

Overall, we have demonstrated that the idea of embedding whole MSAs into a linear representation using standard language modelling approaches can enable the efficient generation of 3-D structures directly from sequence data. This approach enables structural models to be built on a far larger scale than was previously possible. In methodological terms, being able to directly link individual amino acids in a MSA to the outputs of the network, many structural bioinformatics applications could be made easier e.g. modelling variant effects or protein design. At the very least, these direct MSA embedding methods make *de novo* protein structure prediction methods far more efficient and easier to use.

## Acknowledgements

This work was supported by the European Research Council Advanced Grant “ProCovar” (project ID 695558). The authors are grateful to Nicola Bordin and Ian Sillitoe for assistance with the CATH-Gene3D HMM libraries.

## Data availability

The DMPfold2 source code and trained neural network weights are available at https://www.github.com/psipred/DMPfold2 under the terms of the GPLv3 free software licence. Protein structures predicted from the BFD and Pfam are available via the UCL Research Data Repository (doi: 10.5522/04/14979990).

## Supplementary Figures

**Figure S1.**
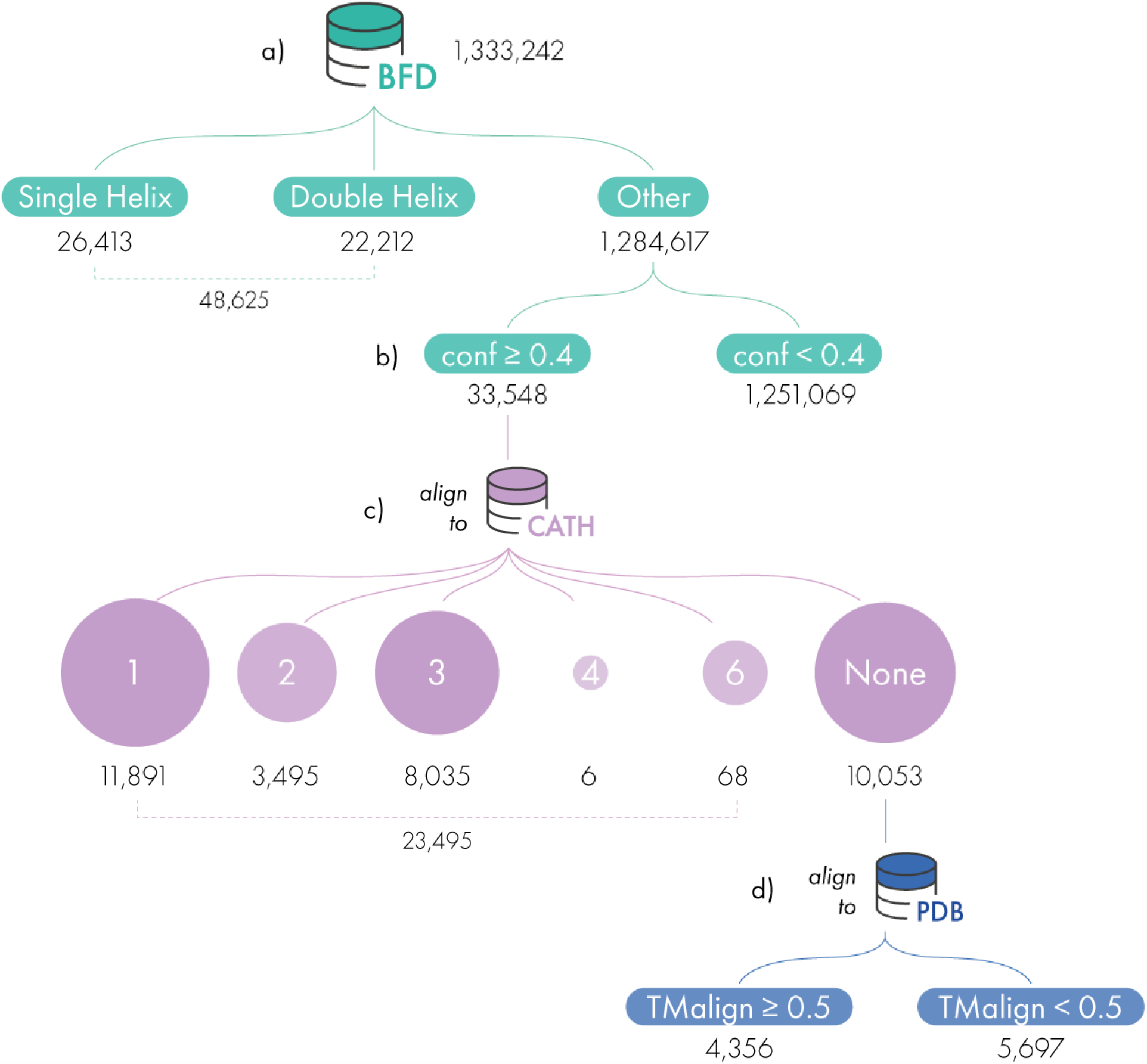
Filtering of BFD models. **a**) Models with folds corresponding to single and double helices were filtered from the ensemble to leave only complex folds. **b**) A confidence threshold of ≥ 0.4 was applied to isolate models likely to be correct. **c**) Each model was searched against known CATH domains. Models which matched CATH domains with a TM-score ≥ 0.5 calculated using TM-align were assigned to the corresponding CATH classes 1-4 and 6. **d**) The remaining ensemble not matched to CATH were searched against a representative set of PDB chains. The number of models at each stage of filtering is represented by the value proximal to each node.

**Figure S2.**
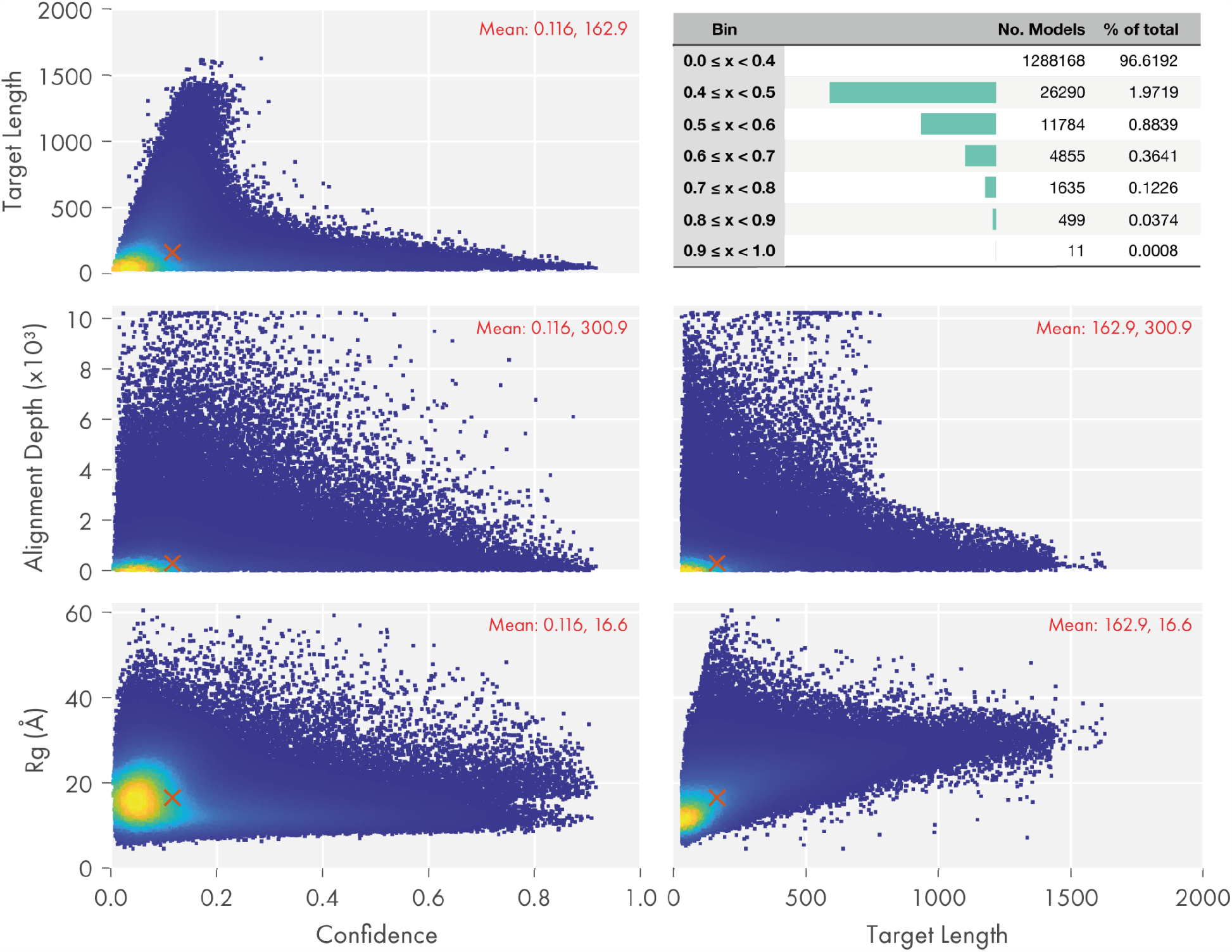
Summary of 1.3 million BFD models. Comparison of target length, alignment depth, radius of gyration (Rg) and model confidence for 1.3 million models. The red ‘x’ marks the mean of each variable, with the numerical values shown in the top right hand corner of each subplot. The table shows the number of models in each confidence bin. Color gradients indicate the density of datapoints, with yellow and dark blue indicating highest and lowest respectively.

**Figure S3.**
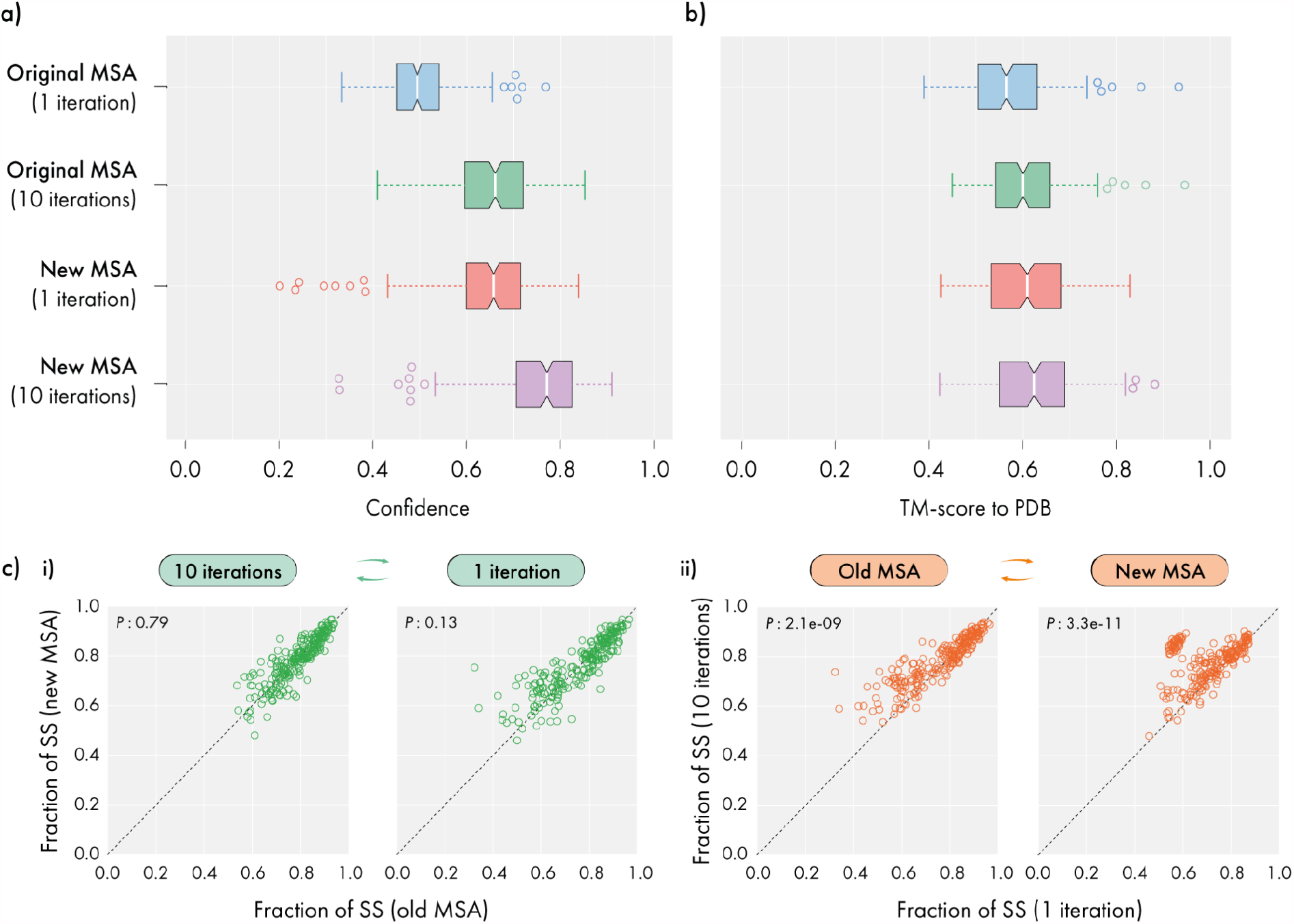
Effect of alignment regeneration and iterative modelling procedure. The models of 225 BFD targets were regenerated to assess the effect of improving the alignment and/or running the end-to-end procedure with 1 or 10 iterations. Box-and-whisker plot of **a**) confidence scores obtained from four different model generation schemes and **b**) highest TM-score using TM-align to PDB structures. **c**) Effect of modelling strategy on model secondary structure content. The fraction of secondary structure was calculated as the total number of residues assigned to the DSSP classes G, H, I, E, B, T or S (i.e. all classes excluding loop/irregular assignments), normalised by the target length. Values in each subplot represent the p-value as determined via a paired t-test.

**Figure S4.**
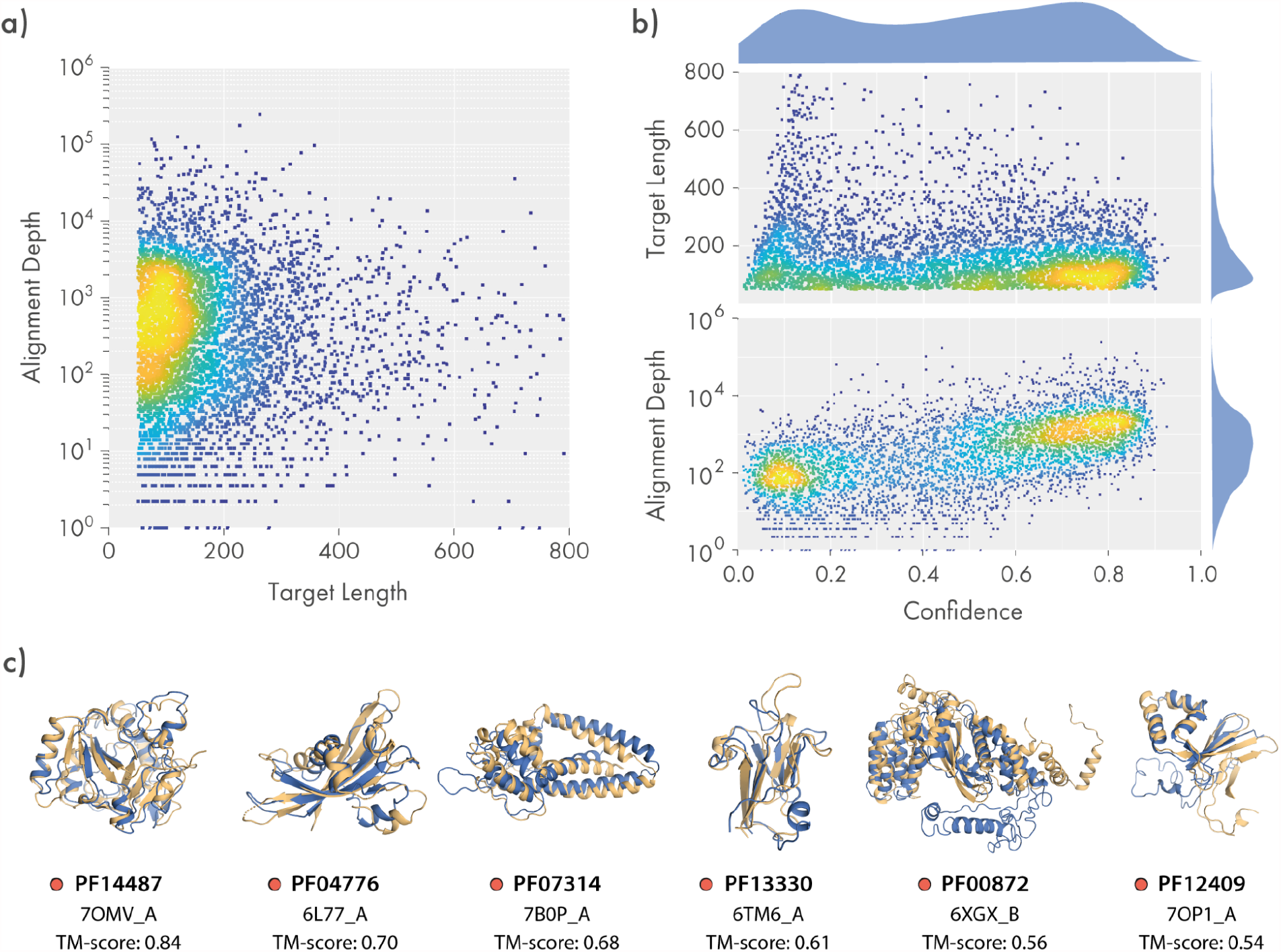
Pfam models generated by DMPfold2. Data represents 4,938 models from Pfam (version 32) generated from alignments used in DMPfold1. **a**) Summary of target length and alignment depth of 4,938 models. **b**) Distributions of model confidence against target length and alignment depth. Of the models generated, 1,988 (40.3%) and 2,950 (59.7%) have a confidence < 0.4 and ≥ 0.4 respectively. Color gradients indicate the density, with yellow and dark blue indicating highest and lowest respectively. **c**) Examples of DMPfold2 models for very hard targets. The mean TM-score across the shown targets are 0.655 for DMPfold2 and 0.564 for C-I-TASSER. Native structures and models are shown in orange and blue respectively.

**Figure S5.**
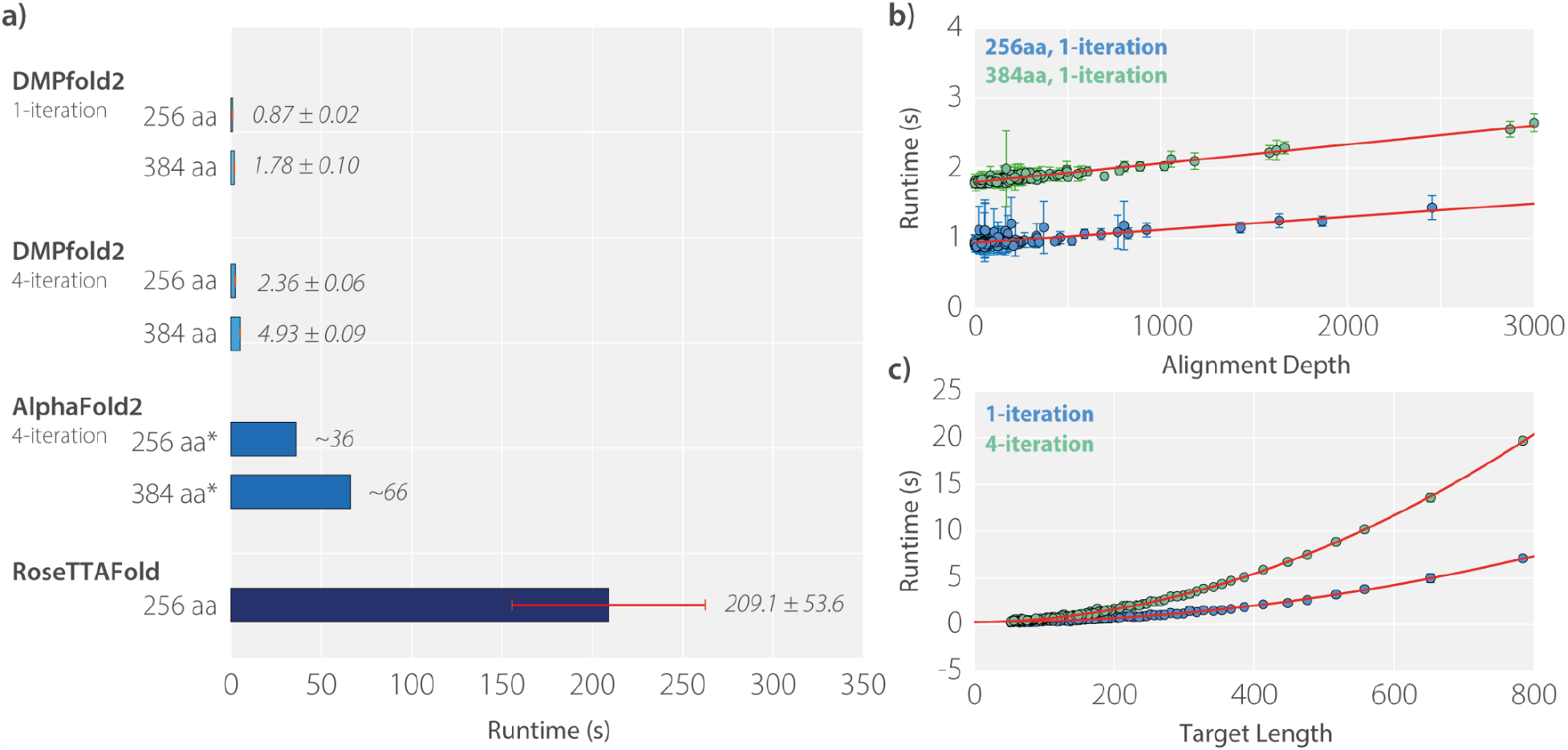
Benchmark of DMPfold2 runtimes. **a**) Comparison of model generation runtimes for DMPfold2 (1 and 4-iteration), AlphaFold2 (4-iteration) and end-to-end RoseTTAFold. Values for DMPfold2 and RoseTTAFold represent the average time taken to generate a model of length 256 and 384 residues. DMPfold2 and RoseTTAfold models were generated using a single NVIDIA GTX 1080 Ti with 11 GB of memory. Timings for the 384-residue target using RoseTTAFold could not be recorded due to memory limitations. AlphaFold2 runtimes (marked by *) were taken directly from its publication and were measured on an NVIDIA V100. **b**) Linear scaling of DMPfold2 runtime with alignment depth. Each datapoint represents a single target of either 256 or 384 residues. **c**) Typical DMPfold2 runtimes for 100 targets of various lengths and alignment depths. Lines indicate quadratic fits of the form *y = ax*^*2*^*+bx+c*. All runtimes were recorded as the average of 10 runs for each target, with error bars indicating standard deviation.

## Supplementary Tables

**Table S1.** Model accuracy on the 39 FM and FM/TBM CASP13 domains, comparing models built using either DMPfold1 or DMPfold2. In the case of DMPfold1, only the top-ranking model selected by each modelling run was evaluated for each target.

| Target | DMPfold1 |  |  | DMPfold2 |  |  |
| --- | --- | --- | --- | --- | --- | --- |
|  | GDT_HA | GDT_TS | TM-Score | GDT_HA | GDT_TS | TM-Score |
| T0949-D1 | 36.434 | 54.845 | 0.626 | 40.310 | 62.984 | 0.720 |
| T0950-D1 | 24.342 | 40.863 | 0.610 | 19.956 | 35.526 | 0.578 |
| T0953s1-D1 | 25.373 | 44.030 | 0.376 | 33.209 | 47.388 | 0.434 |
| T0953s2-D1 | 23.295 | 34.659 | 0.256 | 45.455 | 65.909 | 0.451 |
| T0955-D1 | 41.463 | 52.439 | 0.415 | 56.098 | 74.390 | 0.591 |
| T0957s2-D1 | 36.774 | 58.226 | 0.690 | 31.129 | 51.613 | 0.615 |
| T0958-D1 | 50.974 | 70.779 | 0.704 | 47.727 | 68.506 | 0.668 |
| T0960-D2 | 38.690 | 57.143 | 0.564 | 15.179 | 24.405 | 0.249 |
| T0963-D2 | 41.463 | 59.451 | 0.584 | 15.549 | 26.829 | 0.250 |
| T0968s1-D1 | 46.186 | 67.373 | 0.742 | 41.102 | 62.924 | 0.718 |
| T0968s2-D1 | 41.957 | 63.696 | 0.721 | 41.522 | 63.913 | 0.694 |
| T0969-D1 | 19.986 | 37.641 | 0.633 | 25.989 | 45.056 | 0.697 |
| T0970-D1 | 40.000 | 58.529 | 0.596 | 45.294 | 65.000 | 0.669 |
| T0975-D1 | 19.217 | 33.185 | 0.507 | 24.555 | 43.683 | 0.653 |
| T0978-D1 | 18.765 | 35.472 | 0.584 | 21.065 | 41.223 | 0.662 |
| T0980s1-D1 | 18.750 | 30.288 | 0.310 | 21.875 | 33.654 | 0.356 |
| T0981-D3 | 28.941 | 48.399 | 0.642 | 36.576 | 57.759 | 0.735 |
| T0986s1-D1 | 47.011 | 68.750 | 0.726 | 39.402 | 61.141 | 0.662 |
| T0986s2-D1 | 12.419 | 22.419 | 0.311 | 35.323 | 57.742 | 0.716 |
| T0987-D1 | 22.568 | 40.676 | 0.547 | 31.216 | 53.108 | 0.687 |
| T0987-D2 | 23.597 | 38.903 | 0.520 | 25.893 | 43.750 | 0.576 |
| T0989-D1 | 24.440 | 37.500 | 0.443 | 9.701 | 19.590 | 0.244 |
| T0989-D2 | 16.071 | 29.464 | 0.340 | 11.607 | 23.214 | 0.277 |
| T0990-D1 | 44.408 | 65.461 | 0.611 | 52.961 | 74.013 | 0.745 |
| T0990-D3 | 11.502 | 19.601 | 0.290 | 19.366 | 32.746 | 0.477 |
| T0992-D1 | 50.467 | 72.430 | 0.773 | 48.364 | 69.860 | 0.761 |
| T0997-D1 | 36.216 | 58.243 | 0.722 | 38.378 | 60.811 | 0.761 |
| T0998-D1 | 10.843 | 18.825 | 0.269 | 13.554 | 21.988 | 0.282 |
| T1000-D2 | 27.785 | 46.807 | 0.700 | 28.736 | 49.660 | 0.750 |
| T1001-D1 | 28.237 | 46.763 | 0.560 | 35.072 | 57.374 | 0.672 |
| T1005-D1 | 21.472 | 39.034 | 0.613 | 27.607 | 47.469 | 0.706 |
| T1008-D1 | 19.481 | 27.597 | 0.266 | 24.026 | 37.338 | 0.304 |
| T1010-D1 | 41.190 | 58.810 | 0.711 | 32.024 | 51.190 | 0.672 |
| T1015s1-D1 | 27.273 | 42.898 | 0.452 | 34.659 | 54.261 | 0.559 |
| T1017s2-D1 | 27.600 | 43.400 | 0.502 | 39.800 | 61.800 | 0.733 |
| T1019s1-D1 | 33.621 | 53.879 | 0.433 | 49.569 | 70.259 | 0.652 |
| T1021s3-D1 | 27.410 | 45.783 | 0.576 | 35.843 | 55.271 | 0.664 |
| T1021s3-D2 | 13.918 | 26.031 | 0.287 | 50.258 | 72.423 | 0.775 |
| T1022s1-D1 | 23.878 | 39.904 | 0.508 | 30.449 | 49.199 | 0.594 |
| **Mean** | 29.334 | 45.902 | 0.531 | 32.728 | 51.153 | 0.590 |

**Table S2.** CATH assignment of BFD models. The number of models assigned to each CATH class and not assigned to a CATH class are shown, along with the number of superfamilies present in each class in CATH.

| **Class** | **No. Superfamilies** | **% of Total** | **No. Models Assigned** | **% of Total** |
| --- | --- | --- | --- | --- |
| **1** | 945 | 39.46 | 11,891 | 35.44 |
| **2** | 451 | 18.83 | 3,495 | 10.42 |
| **3** | 962 | 40.17 | 8,035 | 23.95 |
| **4** | 4 | 0.17 | 6 | 0.02 |
| **6** | 32 | 1.34 | 68 | 0.20 |
| **None** |  |  | 10,053 | 29.97 |
| **Total** | 2,395 |  | 33,548 |  |

